# *Vegfa* expression is activated through positive and negative transcriptional regulatory networks controlled by the ETS factor Etv6 *in vivo*

**DOI:** 10.1101/425751

**Authors:** Lei Li, Rossella Rispoli, Roger Patient, Aldo Ciau-Uitz, Catherine Porcher

## Abstract

VEGFA signaling is crucial for physiological and pathological angiogenesis and hematopoiesis. Although many context-dependent signaling pathways downstream of VEGFA have been uncovered, vegfa transcriptional regulation in vivo remains unclear. Here we show that the ETS transcription factor, Etv6, positively regulates vegfa expression during Xenopus blood stem cell development through multiple transcriptional inputs. In agreement with its established repressive functions, Etv6 directly inhibits the expression of the vegfa repressor, foxo3. Surprisingly, it also directly activates the expression of the vegfa activator, klf4. Finally, it indirectly binds to the vegfa promoter where it co-localizes with Klf4. Klf4 deficiency downregulates vegfa expression and significantly decreases Etv6 binding to the vegfa promoter, indicating that Klf4 recruits Etv6 to the vegfa promoter. Thus, our work uncovers a dual function for Etv6, as both a transcriptional repressor and activator, in controlling a major signaling pathway involved in blood and endothelial development in vivo. Given the established relationships between development and cancer, this elaborate gene regulatory network may inform new strategies for the treatment of VEGFA-dependent tumorigenesis.

---

Vascular Endothelial Growth Factor A (VEGFA) signaling is critical for both physiological and pathological processes in the adult, including hematopoiesis, angiogenesis and solid tumor progression. In hematopoiesis, VEGFA regulates hematopoietic stem cell (HSC) survival and function in the bone marrow through both cell intrinsic and extrinsic mechanisms^1,2^. In angiogenesis, VEGFA is required for endothelial cell proliferation, migration and organization in three dimensions to form new vessels during vascular remodeling^3^. Finally, in cancer, VEGFA signaling plays essential roles in the regulation of tumor angiogenesis and metastasis^4^, and has emerged as a key anti-angiogenic target in a wide variety of cancer therapies^5^. However, inhibitors of VEGFA-mediated signaling often trigger limited responses in both human tumors and mouse models of cancer, due to the development of evasive and intrinsic resistance mechanisms^5^.

VEGFA signaling is also essential during embryogenesis where it plays pivotal roles in the development of the endothelial and hematopoietic systems. Knockout of VegfA or VegfA receptor (Vegfr1, 2 or 3) genes in mice results in early lethality owing to severe defects in vascular development^6^. Mice deficient in Vegfr2 (also known as Flk1 or Kdr) show an absence of yolk sac blood islands and reduction of CD34^+^ hematopoietic progenitors^7^. In vitro, VEGFA is required for the specification of FLK1^++^ mesoderm into hematopoietic and cardiovascular lineages in mouse embryonic stem cell differentiation models^8,9^. In embryos, vegfa is essential for HSCs emergence and has been shown to be required at several stages of their programming^10–12^. In Xenopus, acting in a paracrine manner from the somites, Vegfa is required for the establishment of definitive hemangioblasts (DHs, precursors of the dorsal aorta (DA) and HSCs) in the dorsal lateral plate (DLP) mesoderm^11,12^. Later, Vegfa secreted by the hypochord guides the migration of DA precursors from the DLP to the midline^11,12^.

Although critical to understand the development of the vascular and hematopoietic systems, very little is known about the regulation of the complex spatio-temporal expression pattern of vegfa during embryogenesis. Because developmental and tumour biology and, in particular, angiogenic and metastatic processes share signaling and transcriptional pathways^13^, a better understanding of the regulatory networks lying upstream of VEGFA during specification of the hemato-vascular lineage may help identify new druggable targets relevant to VEGFA-dependent diseases.

ETV6, a transcriptional repressor belonging to the ETS family of transcription factors (TFs)^14–16^, is a key regulator of hematopoietic, angiogenic and tumorigenic processes, and has been linked to vegfa regulation. It is essential for bone marrow hematopoiesis^17,18^ and is involved in many chromosomal rearrangements which lead to childhood leukaemias^19^. Additionally, germline and somatic mutations in ETV6 have recently been shown to be involved in other malignancies such as skin, colon, mammary and salivary gland cancers^20–22^. In mouse embryos, Etv6 homozygous deletion leads to early embryonic death because of defective yolk sac angiogenesis^23^.

Previously, we showed that Etv6 specifies DHs and HSCs in developing Xenopus embryos through positive regulation of vegfa expression in the somites at stage 22 of development^12^. Because Etv6 is considered a transcriptional repressor^14,15,24^, this suggested an indirect regulatory mechanism of action whereby Etv6 directly represses expression of a vegfa transcriptional repressor. To test this hypothesis, we set out to investigate the mechanism by which Etv6 activates vegfa expression in the somites. Using genome-wide and functional analyses, we identify Etv6 direct target genes and show that Etv6 employs multiple regulatory mechanisms. To allow vegfa expression, Etv6 directly represses foxo3, a known transcriptional repressor of VegfA in cancer cells^25,26^, thus participating in a double negative gate. Unexpectedly, Etv6 also directly activates expression of klf4, a known activator of VegfA in endothelial cells and a regulator of metastatic processes^27,28^. Consistent with a feed-forward loop mechanism, Etv6 also binds the vegfa promoter. This recruitment is mediated by Klf4, suggesting co-operation between the two proteins to activate vegfa expression. Thus, our work demonstrates that vegfa expression in the somites is regulated by a complex gene regulatory network (GRN) with Etv6 acting both as a repressor and, surprisingly, an activator. These findings further our understanding of the in vivo regulation of vegfa and provide a platform for the study of the processes underlying VEGFA-dependent tumorigenesis, such as those involving FOXO3 and KLF4, and for the establishment of novel therapeutic strategies.

## Results

### Etv6 genome-wide occupancy in the somites of Xenopus embryos

To investigate the mechanism by which Etv6 regulates vegfa expression in the somites, we first set out to identify Etv6 genomic targets through ChIP-seq (Figure 1a). To this end, we generated ChIP-grade anti-Xenopus-Etv6 polyclonal antibodies in rabbits. Three short peptides were designed (Supplementary Figure 1a) and used to generate six polyclonal antibodies (two/peptide). Four of these antibodies detected Etv6 protein on Western blots (Supplementary Figure 1b) and were tested for their ability to immunoprecipitate endogenous Etv6 protein from protein extracts generated from the somites of stage 22 embryos. Two antibodies (Etv6-2a and Etv6-2b) immunoprecipitated endogenous Etv6 protein (Supplementary Figure 1c). Etv6-2a showed greater affinity to Etv6 and, therefore, was selected for ChIP-seq experiments. Chromatin immunoprecipitation was performed on three independent biological samples obtained from the somites of wild-type (WT) stage 22 embryos (Figure 1a). Indexed libraries were generated from immunoprecipitated DNA and control input, samples were pooled and sequenced, and reads mapped to the X. laevis genome.

**Figure 1.**
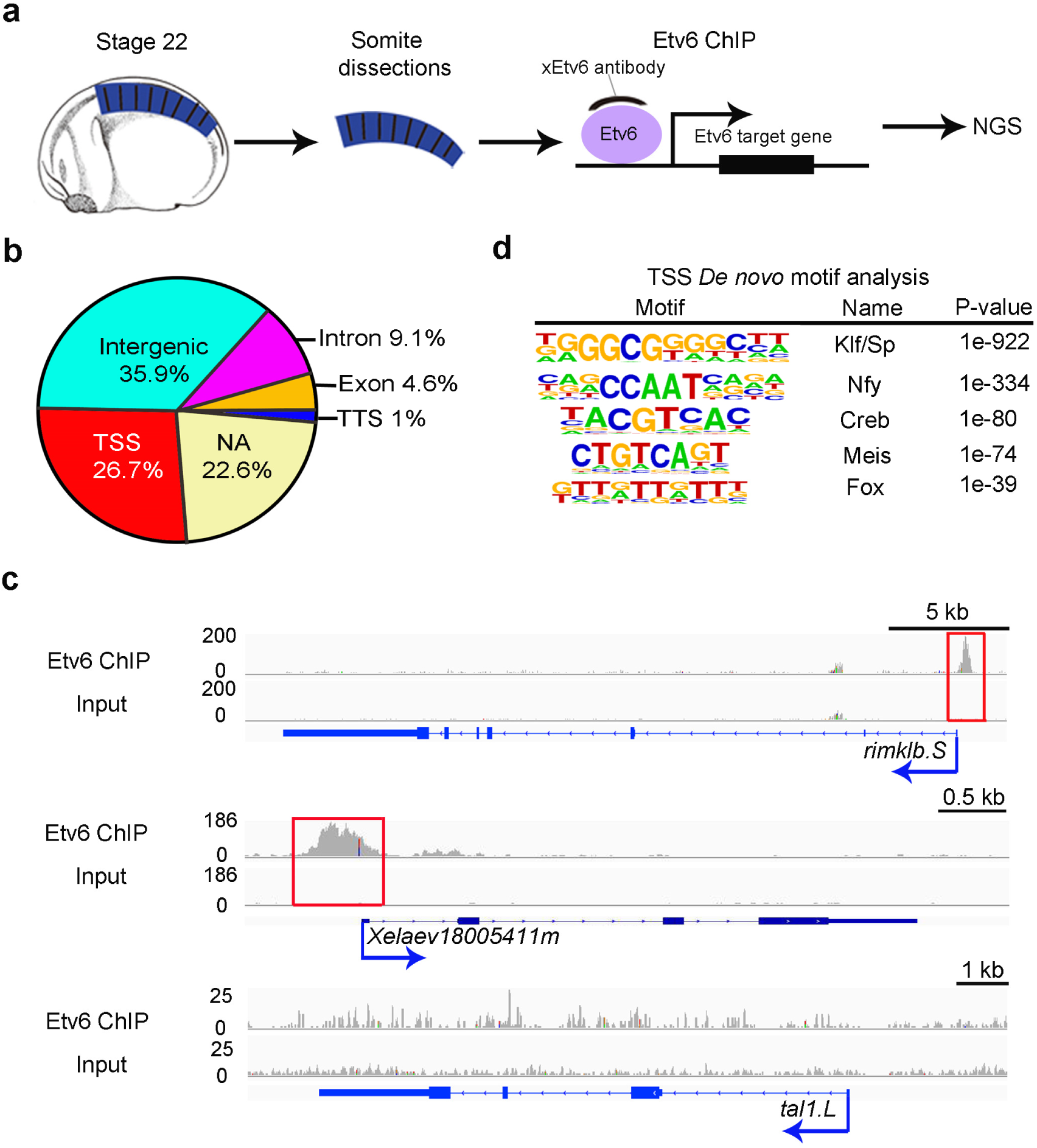
Genome-wide occupancy of Etv6 in *Xenopus* somites at stage 22. (**a**) Experimental design of Etv6 ChIP-seq assay. The somites were dissected from stage 22 Xenopus laevis embryos, homogenized and fixed, and subjected to Etv6 ChIP-seq. NGS, next generation sequencing. (**b**) Genomic distribution of the 9,128 Etv6 ChIP-seq peaks throughout the X. laevis genome. NA, not annotated; TSS, transcription start site; TTS, transcription termination site. (**c**) Integrative genome viewer (IGV) showing examples of Etv6 ChIP-seq tracks. Etv6 binding is enriched in the TSS region of rimklb.S and Xelaev18005411m (red boxes) but not in the TSS region of tal1.L. Input is shown as control. (**d**) De novo motif analysis of ETV6 peaks located in the TSS region. Five out of the 22 overrepresented motifs are shown.

9,128 Etv6 consistent peaks across the three biological replicates were identified (Supplementary Table 1). The genomic distribution of the peaks revealed Etv6 occupancy over distinct features including transcription start sites (TSS), transcription termination sites (TTS), exons, introns and intergenic regions (Figure 1b). Additionally, 22.6% of the peaks mapped to genomic regions not yet annotated (NA). For all subsequent analyses, and as a first step to identify Etv6 direct transcriptional targets, we focused on the 2,440 peaks (26.7%) located in TSS regions that could confidently be associated with 2416 genes. In Figure 1c, we show the profiles of two genes with the highest enrichment of Etv6 in their TSS region (rimklb and Xelaev18005411m), and of tal1, a hematopoietic marker gene which is not expressed in the somites at stage 22^11^ and devoid of Etv6 peaks at its TSS region.

Next, we performed de novo motif analysis on the sequences underlying all Etv6 peaks associated with TSS regions. Etv6 peaks showed significant enrichment of consensus binding sequences corresponding to known TFs (Figure 1d). Surprisingly, ETS family binding motifs were not represented (Supplementary Table 2). Instead, the most over-represented motif belonged to the Klf/Sp family (P value=1e-922), followed by consensus motifs for Nfy, Creb, Meis and Fox TFs. This suggested that, in TSS regions, Etv6 may be preferentially recruited to chromatin indirectly, through interaction with other DNA-binding TFs such as members of the Klf/Sp family.

### Establishing the transcriptome regulated by Etv6 in the somites

To determine the transcriptional targets of Etv6 in the somites of stage 22 embryos, we compared the transcriptome of WT and etv6-deficient somites, the latter generated with a previously validated etv6 antisense morpholino oligonucleotide (MO)^12^ (Figure 2a). Spearman correlation analysis and Principal Component Analysis (PCA) revealed that etv6 deficiency causes significant changes in the transcriptome of the somites (Figure 2b, Supplementary Figure 2). Indeed, differential expression analysis identified a total of 5,186 differentially expressed genes (DEGs, FDR<0.05) with 2,236 genes normally repressed by Etv6 (upregulated in etv6-deficient somites) and 2,950 genes normally activated by Etv6 (downregulated in etv6-deficient somites) (Figure 2c, Supplementary Table 3). Interestingly, gene ontology analysis indicated an enrichment in categories related to positive and negative regulation of transcription (Supplementary Table 4), confirming our hypothesis that Etv6 controls biological processes through the regulation of the expression of transcriptional regulators.

**Figure 2.**
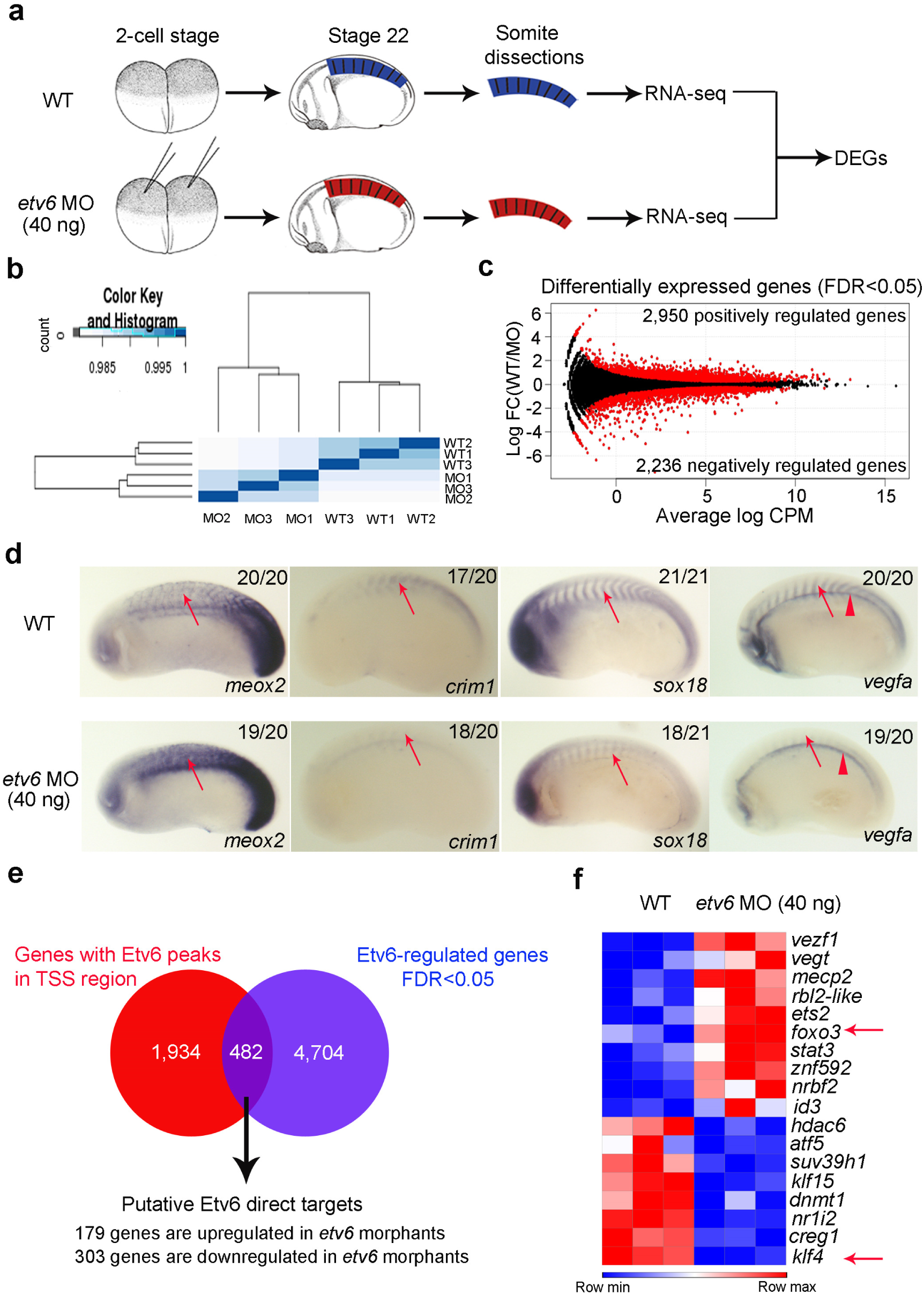
Identification of Etv6 direct transcriptional targets in the somites of stage 22 *Xenopus* embryos. (**a**) Experimental design. RNA-seq was performed on wild type (WT) and etv6-deficient (etv6 MO injected embryos) somite explants dissected from stage 22 embryos. Differentially expressed genes (DEGs) were identified by comparing the transcriptome of these tissues. (**b)** Spearman correlation analysis on triple biological RNA-seq replicates. (**c)** Smear plot showing the fold-change in gene expression of all genes in WT versus etv6-deficient somites (log_2_FC) compared to their expression levels (log_2_CPM). Black dots, non-significant change; red dots, differentially expressed genes (DEGs). The number of positively and negatively regulated DEGs is indicated, (**d)** WISH showing the expression of DEGs in stage 22 WT and etv6-deficient embryos. Meox2 expression is upregulated in the somites (arrows) in etv6-deficient embryos whereas expression of crim1, sox18 and vegfa is downregulated. Note that vegfa expression in the hypochord is unaffected (arrowheads). Embryos are shown in lateral view with anterior to the left and dorsal to the top. Numbers in top right corner indicate the number of embryos exhibiting the phenotype pictured, (**e**) The intersection between DEGs and genes harboring Etv6 ChIP-seq peaks in their TSS region reveals 482 putative direct target genes. (**f)** Foxo3 and klf4 (arrows), known transcriptional regulators of vegfa, are amongst the 18 TFs and chromatin modifiers identified as potential direct transcriptional targets of Etv6.

To validate the RNA-seq data, we next confirmed the differential expression of selected genes in WT and etv6-deficient embryos by Whole-mount in situ hybridization (WISH) (Figure 2d). During vertebrate development, the homeobox genes meox1 and meox2 regulate somitogenesis and myogenesis^29^. In zebrafish, meox1 also represses the expansion of somite-derived endothelial cells thereby limiting HSC development in the DA^30^. The single Xenopus meox gene is repressed by Etv6 in somites (WT/etv6 MO log_2_FC=-0.89 for meox2.L and −0.53 for meox2.S) and WISH confirmed higher expression in etv6-deficient somites (Figure 2d). Crim1, a regulator of VEGFA autocrine signaling in endothelial cells^31^, is positively regulated by Etv6 (WT/etv6 MO log_2_FC =0.68). WISH analysis confirmed down-regulation of crim1 expression in the somites of etv6-deficient embryos (Figure 2d). Sox18, a TF required for the development of blood vessels^32^, is positively regulated by Etv6 in the somites, as revealed by RNA-seq and confirmed by WISH (WT/etv6 MO log_2_FC=1.01, and decreased expression in the somites of etv6-deficient embryos, Figure 2d). Finally, we have previously reported that vegfa expression in the somites is activated by Etv6^12^. Intriguingly, vegfa was not identified as a differentially expressed gene by RNA-seq. WISH analysis did, however, confirm that vegfa is absent in the somites of etv6-deficient embryos. It also revealed strong vegfa expression in the hypochord of both WT and etv6-deficient embryos (Figure 2d), thus providing an explanation for the absence of vegfa from the DEG list, as the explants used for RNA extraction contained the hypochord, the unperturbed expression of vegfa in the etv6-deficient hypochord masked its down-regulation in the somites. More importantly, this demonstrates that Etv6 specifically controls expression of vegfa in the somites. In conclusion, our RNA-seq data reveals a robust transcriptional response to etv6 knock-down.

### Identification of Etv6 direct target genes

To identify Etv6 direct transcriptional targets, we compared the list of genes harboring Etv6 peaks in their TSS regions (2,416 genes) to the list of DEGs (5,186 genes). This revealed 482 putative direct target genes with 498 peaks in their TSS regions (Figure 2e, Supplementary Table 5). Surprisingly, the majority of Etv6 target genes (303/482, 63%) were downregulated in the somites of etv6-deficient embryos, strongly indicating that Etv6, typically considered as a transcriptional repressor^19^, acts mainly as an activator of gene expression in the somites at stage 22. De novo motif analysis on the peaks linked to genes repressed by Etv6 identified Klf/Sp, Nfy, Creb and Ets motifs, whilst peaks linked to genes activated by Etv6 contained Klf/Sp, Nfy, Creb and Hox motifs, and were not enriched in ETS-binding cis-elements (Supplementary Table 6). Altogether, this strongly suggested that recruitment of Etv6 to genes it normally activates relies predominantly on other DNA-binding TFs.

To identify the transcriptional regulators that could mediate Etv6 control of vegfa expression, we next focused on the 18 TFs and chromatin modifiers detected amongst the Etv6 putative direct targets with fold-change in gene expression >1.5 (Figure 2f). Many of these transcriptional regulators are involved in the control of vegfa expression in higher vertebrates^33-37^. Amongst those, foxo3 (Forkhead box o3), a transcriptional repressor of vegfa in cancer cells^25,26^, and klf4 (Krüppel-like factor 4), a transcriptional activator of vegfa in endothelial cells and a regulator of metastatic processes^27,28^, were respectively repressed and activated by Etv6 (Figure 2f). We hypothesized that Foxo3 and Klf4 may contribute to the regulation of vegfa expression in the somites.

### Etv6 prevents Foxo3-mediated repression of vegfa in the somites

To investigate whether Etv6 could be regulating vegfa expression by repressing foxo3 expression, we first confirmed binding of Etv6 to both foxo3 genes in the X. laevis genome. X. laevis is an allotetraploid organism with a genome organized into two homologous sub-genomes, referred to as L and S, and most genes (>56%) are represented by two distinct homologous copies which can be regulated differently^38^. ChIP-seq tracks in Figure 3a show that Etv6 peaks are present in the TSS region of both foxo3.L and foxo3.S and Etv6 binding in these regions was confirmed by ChIP-qPCR (Figure 3b). These peaks contained one Ets (5’-G(A/T)GGAAG(G/T)-3’) and several Klf/Sp binding motifs. Furthermore, upregulation of foxo3 in the somites of etv6-deficient embryos was confirmed by WISH with probes designed to target both foxo3.L and foxo3.S (Figure 3c) and RT-qPCR on RNA extracted from stage 22 somites with primers designed to detect both foxo3 genes (Figure 3d). In conclusion, both foxo3.L and foxo3.S are directly repressed by Etv6 in the somites.

**Figure 3.**
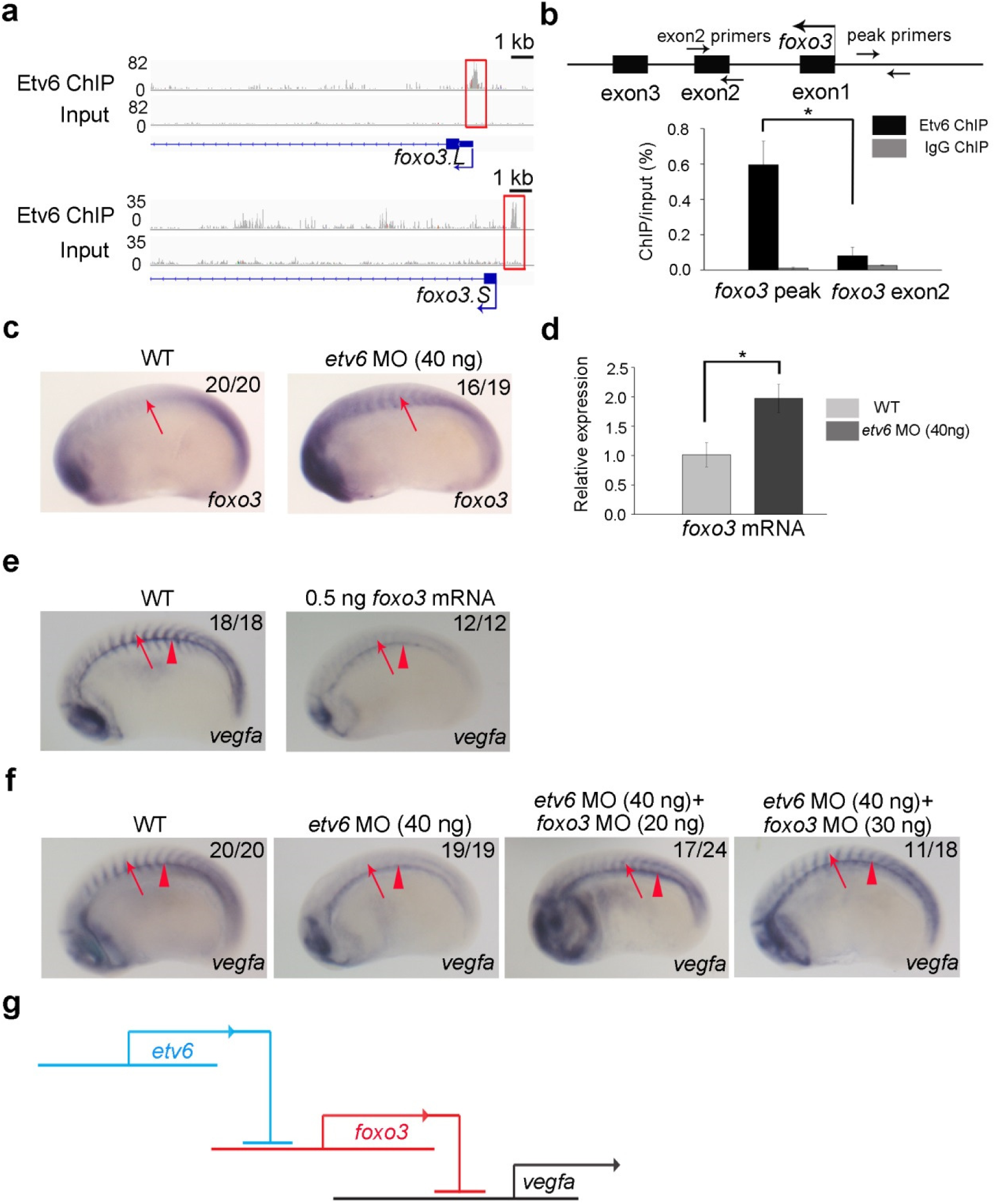
Etv6 prevents Foxo3-mediated repression of *vegfa* in the somites. (**a**) IGV showing Etv6 peaks (red boxes) in the TSS region of foxo3.L and foxo3.S. Input is shown as control. (**b**) ChIP-qPCR analysis confirming that Etv6 binding is enriched in the foxo3 promoter region. The diagram above the histogram depicts the foxo3 locus and the location of the primers used to amplify the region of the Etv6 peak in the foxo3 TSS region (peak primers) and a negative control region in exon2 (exon2 primers). Primers were designed to target both foxo3.L and foxo3.S. IgG ChIP was used as negative control. Error bars represent SEM of three biological replicates. *P=0.014, two-tailed Student’s t-test. (**c**) WISH showing that foxo3 expression is upregulated in the somites (arrows) of etv6-deficient embryos. (**d**) RT-qPCR confirming that foxo3 is upregulated in stage 22 etv6-deficient somites. Expression was normalized to odc1. Error bars represent SEM of three biological replicates. *P=0.020, two-tailed Student’s test. (**e**) WISH showing that overexpression of foxo3 (0.25 ng foxo3.L mRNA+0.25 ng foxo3.S mRNA) blocks vegfa expression in the somites (arrows) whereas expression in the hypochord (arrowheads) is unaffected. (**f**) WISH showing that blocking foxo3 translation with MOs rescues vegfa expression in the somites (arrows) of etv6-deficient embryos. Arrowheads indicate expression in the hypochord. (**g**) Diagram illustrating that vegfa expression in the somites requires the repression of foxo3 by Etv6, i.e. Etv6 represses a repressor of vegfa. Images in **c**, **e** and **f** show stage 22 embryos in lateral view with anterior to the left and dorsal to the top. Numbers in top right corner indicate the number of embryos exhibiting the phenotype pictured.

Next, we wanted to test the hypothesis that Foxo3 may repress the expression of vegfa in the somites. First, we determined whether upregulation of foxo3 was sufficient to repress vegfa in the somites by injecting foxo3 mRNA into 2-cell stage embryos and testing vegfa expression by WISH at stage 22. Indeed, vegfa expression was dramatically downregulated in the somites of embryos injected with exogenous foxo3 mRNA (Figure 3e) whereas expression in the hypochord was unaffected. This indicated that foxo3, like Etv6, specifically regulates the expression of vegfa in the somites, supporting the notion that Etv6 positively regulates the expression of vegfa through repression of foxo3. To further confirm that vegfa downregulation in the somites of Etv6-deficient embryos is mediated by Foxo3, we designed MOs that efficiently block the translation of foxo3 (Supplementary Figure 3a-b). Consistent with the absence of foxo3 expression in the somites of stage 22 embryos (Figure 3c and Supplementary Figure 4), injection of foxo3 MOs had no effect on vegfa expression in the somites of WT embryos (Supplementary Figure 3c). However, co-injection of foxo3 MOs with etv6 MOs rescued the expression of vegfa in the somites of etv6-deficient embryos (Figure 3f), indicating that Etv6 prevents Foxo3-mediated repression of vegfa in the somites (Figure 3g). In cancer cells, FOXO3 has been reported to repress vegfa expression by directly binding to the vegfa promoter region^25,26^. We have identified a putative Foxo3 binding site (5′-GTAAACA-3′)^39^ in the promoter region of Xenopus vegfa using Jaspar^40^, strongly suggesting that Foxo3 can bind the vegfa promoter directly and repress vegfa expression in the somites of etv6-deficient embryos.

### Etv6 acts as a direct transcriptional activator of klf4

The klf4 gene is a candidate direct target of Etv6 in the somites (Figure 2f). Interestingly, KLF4 has been shown in human retinal microvascular endothelial cells (HRMECs)^27^ and human umbilical vein endothelial cells (HUVECs)^28^ to activate vegfa expression through direct binding to its promoter. We therefore hypothesized that Etv6 could activate vegfa expression in the somites through Klf4.

We first validated binding of Etv6 in the promoter regions of the two Xenopus klf4 genes, klf4.L and klf4.S (Figure 4a-b), and confirmed by WISH (Figure 4c) and RT-qPCR (Figure 4d) on stage 22 etv6-deficient embryos that expression of klf4 in the somites was dependent on Etv6. Additionally, we demonstrated by western blot that Klf4 protein levels were dramatically reduced in the somites of etv6-deficient embryos (Figure 4e, etv6 MO). Taken together, this suggested that klf4 expression in the somites is directly activated by Etv6.

**Figure 4.**
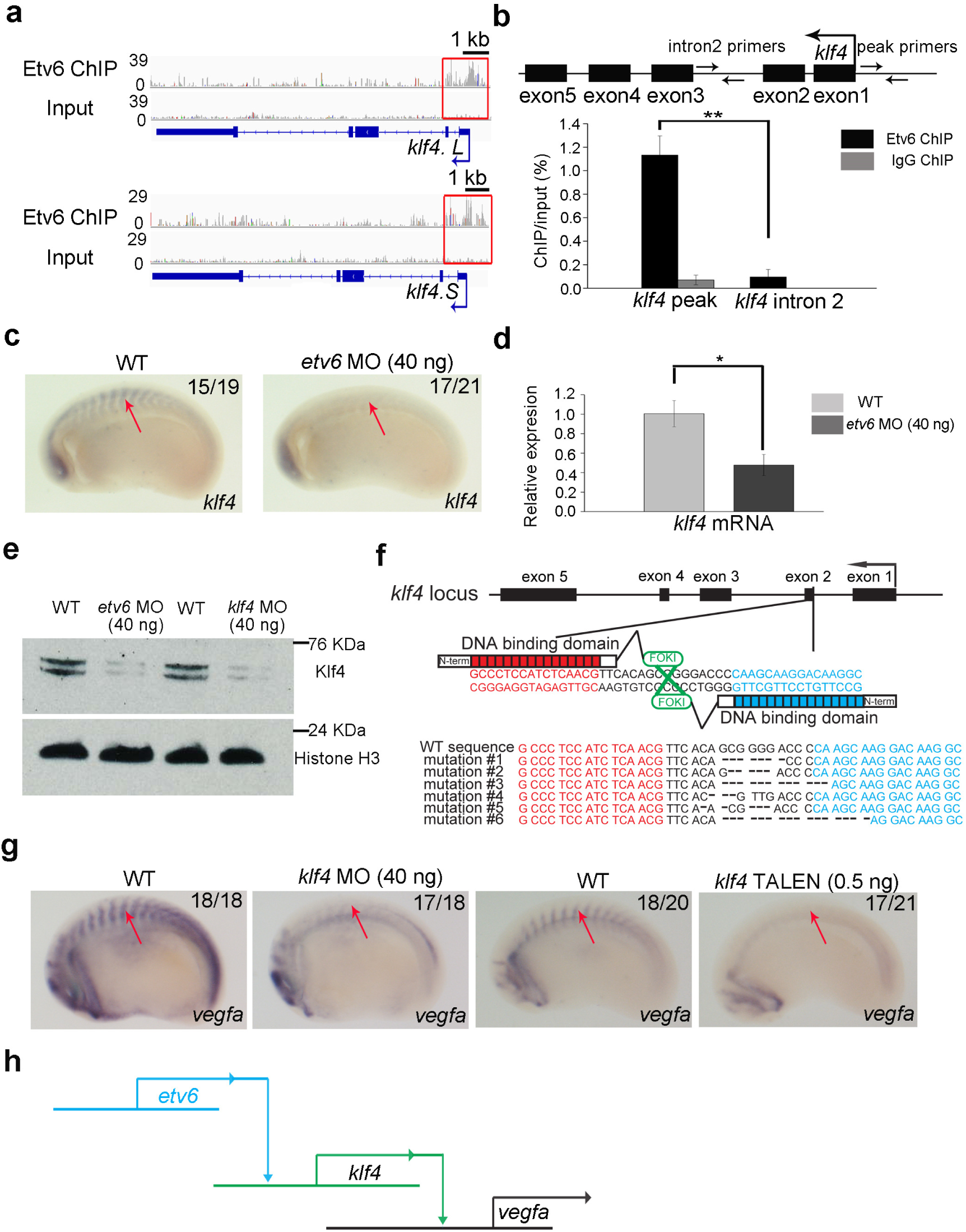
Etv6 positively regulates *vegfa* expression through transcriptional activation of *klf4*. (**a**) IGV showing Etv6 peaks (red boxes) in the TSS region of klf4.L and klf4.S. Input is shown as control. (**b**) ChIP-qPCR analysis confirming that Etv6 is enriched in the klf4 promoter region. The diagram above the histogram depicts the klf4 locus and the location of the primers used to amplify the region of the Etv6 peak in the klf4 TSS region (peak primers) and a negative control region in intron2 (intron2 primers). Primers were designed to target both klf4.L and klf4.S. IgG ChIP was used as negative control. Error bars represent SEM of three biological replicates. **P=0.0036, two-tailed Student’s t-test. (**c**) WISH showing that klf4 expression in the somites (arrows) is downregulated in etv6-deficient embryos. (**d**) RT-qPCR confirming that klf4 is downregulated in stage 22 etv6-deficient somites. Expression was normalized to odc1. Error bars represent SEM of three biological replicates. *P=0.020, two tailed Student’s test. (**e**) Western blot showing that Klf4 protein is depleted in etv6-deficient embryos and that klf4 MO blocks efficiently the translation of Klf4. Histone H3 was used as a loading control. (**f**) Klf4 TALEN design. Top, diagram showing the sequences in klf4 exon 2 targeted by the TALENs; bottom, DNA alignment showing the range of mutations generated by TALEN activity. Genomic DNA was obtained from stage 22 WT and TALEN-injected somites, and subjected to Sanger sequencing. TALEN-injection caused mutations in 78% of the clones sequenced. (**g**) WISH showing that vegfa expression in the somites (arrows) is downregulated in both klf4 MO- and klf4 TALEN-injected embryos. (**h**) Diagram illustrating that Etv6 positively regulates vegfa expression in the somites through transcriptional activation of klf4, a transcriptional activator of vegfa.

Analysis of the expression pattern of klf4 during early embryogenesis indicates that its expression in the somites is remarkably transient as it is only detected in this tissue at stage 22 (Supplementary Figure 5). Importantly, vegfa expression in the somites is absolutely dependent on Etv6 at this stage^11^. This is also the stage when vegfa secreted from the somites is required for the programming of definitive hemangioblasts in the lateral plate mesoderm^12^. In order to investigate the role of Klf4 in vegfa expression and HSC programming, we designed a MO which efficiently blocks the in vivo translation of Klf4 (Figure 4e, Klf4 MO) as well as TALENs (Transcription activator-like effector nucleases) which efficiently generate null mutations in the second exon of klf4 (Figure 4f). Klf4-deficient embryos generated either by MO or TALEN injection showed a dramatic downregulation of vegfa expression in the somites at stage 22 (Figure 4g). Thus, Klf4 is required for vegfa expression in the somites. Furthermore, definitive hemangioblast specification in the lateral plate mesoderm, as indicated by tal1 expression at stage 26-28, and hemogenic endothelium emergence in the dorsal aorta, as indicated by runx1 expression at stage 39, failed in klf4-deficient embryos (Supplementary Figure 6). Therefore, Klf4 is essential for vegfa expression in the somites and the programming of HSCs during embryogenesis, thus recapitulating the Etv6 functions.

Collectively, our studies show that Etv6 positively regulates the expression of vegfa in the somites through direct transcriptional activation of klf4, a transcriptional activator of vegfa (Figure 4h).

### Klf4 is required for Etv6 binding to the vegfa promoter

Analysis of Etv6 ChIP-seq peaks associated with the vegfa genes revealed binding of Etv6 in the vegfa.L (Figure 5a), but not vegfa.S (data not shown), promoter region (−552 to −244 bp upstream of vegfa.L TSS). Analysis of the genomic structure of vegfa.S indicated that it is a gene undergoing degeneration and it is not transcribed during embryogenesis (Xenbase expression data after Sessions et al^38^). Thus, only one vegfa gene, vegfa.L, is functional in X. laevis (referred to as vegfa hereafter) and bound by Etv6 at its TSS. To test whether vegfa could be directly bound by Etv6, we performed Etv6 ChIP-PCR on the vegfa promoter and confirmed the enrichment of Etv6 in this region (Figure 5b-c). However, we did not find ETS binding motif under this peak, suggesting that Etv6 cannot directly bind to the vegfa promoter. Instead, two binding motifs for the Klf/Sp family with high matrix scores were identified (Supplementary Table 7). This is consistent with our de novo motif analysis showing that Klf/Sp binding motifs are the most overrepresented sequences and that genes directly activated by Etv6 do not have ETS motif enrichment under Etv6 peaks in TSS regions.

**Figure 5.**
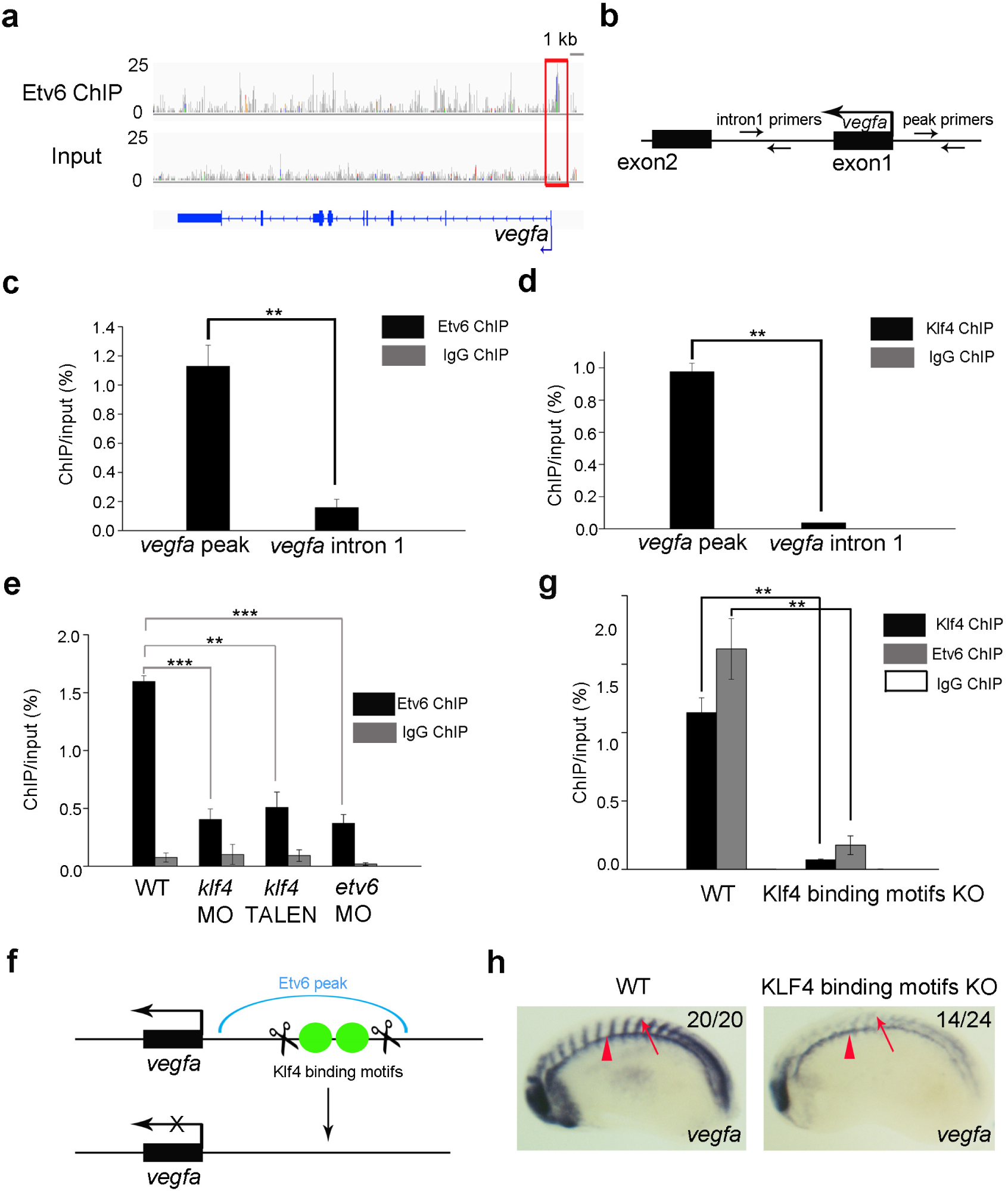
Klf4 is required for the recruitment of Etv6 to the *vegfa* promoter. (**a**) IGV view of Etv6 peaks reveals a peak (red box) in the TSS region of vegfa. Input is shown as control. (**b**) Partial representation of the vegfa locus showing the location of the primers used to amplify the region of the Etv6 peak in the vegfa TSS region (peak primers) and a negative control region in intronl (intronl primers). (**c-d**) ChIP-qPCR analysis on stage 22 somite explants confirming that Etv6 (**c**) and Klf4 (**d**) are enriched in the promoter region of vegfa. IgG ChIP was used as negative control. Error bars represent SEM of three biological replicates. **P=0.003 (**c**), **P=0.0011 (**d**), two-tailed Student’s t-test. (**e**) The enrichment of Etv6 in the vegfa promoter is significantly reduced in klf4-depleted (40 ng klf4 MO and 0.5 ng klf4 TALEN) stage 22 somites. This reduction is comparable to that produced by the depletion of Etv6 (40 ng etv6 MO), strongly indicating that Klf4 is required for Etv6 recruitment to the vegfa promoter. IgG ChIP was used as negative control. Error bars represent SEM of three biological replicates. ***P=0.0002 WT vs klf4 MO, **P=0.002 WT vs klf4 TALEN, ***P=0.000065 WT vs etv6 MO, two-tailed Student’s t-test. (**f**) Diagram showing the TALENs designed to delete the klf4 binding motifs under the Etv6 peak in the vegfa promoter region. (**g**) The enrichment of Klf4 and Etv6 in the vegfa promoter region is significantly reduced in the somites of Klf4 binding motifs knockout (KO) embryos. IgG ChIP was used as negative control. Error bars represent SEM of three biological replicates. **P=0.0033 for Klf4 ChIP-qPCR, **P=0.0047 for Etv6 ChIP-qPCR, two-tailed Student’s t-test. (**h**) WISH showing that vegfa expression is downregulated in the somites of Klf4 binding motifs KO embryos, whereas expression in the hypochord (arrowheads) is not affected.

As Klf4 is required for vegfa expression in the somites, we wondered whether it was involved in Etv6 binding to the vegfa promoter. Therefore, we performed Klf4 ChIP-qPCR on the vegfa promoter using the same primers as for Etv6 ChIP-qPCR (Figure 5b-c). This confirmed that Klf4 binding is indeed enriched in the promoter of vegfa and co-localises with Etv6 binding (Figure 5d), supporting the hypothesis that Klf4 is involved in Etv6 recruitment to the vegfa promoter. To further confirm this notion, Etv6 ChIP-qPCR was carried out on klf4-deficient somites (Figure 5e). Etv6 enrichment in the vegfa promoter was severely reduced in klf4-deficient embryos, a reduction similar to that observed in etv6-deficient embryos (Figure 5e). Importantly, etv6 expression was unaffected in the somites of klf4-depleted embryos (Supplementary Figure 7). Therefore, Klf4 is required for Etv6 recruitment to the vegfa promoter.

Finally, to functionally test the role of the two Klf4 binding motifs identified under the Etv6 peak in the vegfa promoter, we designed TALENs that efficiently deleted them (Figure 5f, Supplementary Figure 8). In the somites of stage 22 embryos with Klf4 binding motifs deleted, the enrichment of Klf4 and Etv6 in the vegfa promoter region was dramatically decreased (Figure 5g). Importantly, vegfa expression in the somites was downregulated in embryos deleted for Klf4 binding motifs, whereas expression in the hypochord was not affected (Figure 5h), strongly indicating that these Klf4 binding sites are specifically required for vegfa expression in the somites of stage 22 embryos.

Taken together, our results show that klf4 is not only a direct target of Etv6, but also acts as its recruiting factor to the vegfa promoter.

## Discussion

In this study, we show that expression of the important signaling molecule vegfa is under tight transcriptional control by the ETS TF Etv6 during the early stages of hematovascular development. Previously, we showed that Etv6 is essential for expression of vegfa in the somites^12^. We now demonstrate that it works through both repressive and activating transcriptional mechanisms. As discussed below, this reveals unexpected molecular mechanisms engaged by Etv6 and a complex gene regulatory network upstream of vegfa in vivo.

To establish how Etv6 controls vegfa expression in the somites, we identified Etv6 direct target genes by focusing on genes that are both abnormally expressed in etv6 deficient somites and bound by Etv6 in their TSS. This stringent approach revealed numerous biologically relevant targets. Indeed, many of the TFs and chromatin modifiers identified amongst Etv6’s 482 direct target genes have previously been directly or indirectly implicated in the regulation of VegfA in endothelial and cancer cells. In addition to Foxo3 and Klf4 (the two targets examined in this study), these are Ets2, shown to down-regulate VegfA expression in human umbilical vein endothelial cells (HUVECs) treated with arsenic trioxide (ATO)^37^, the tumor suppressor Rbl2 (also known as RB2/p130), that inhibits tumor formation in mice by inhibiting VegfA-mediated angiogenesis^35^, the transcriptional repressor Mecp2, that represses VegfA expression by binding to methylated CpG islands in the VegfA promoter^33^, and the transcriptional regulator Creg1, that regulates human endothelial homeostasis in vivo and promotes vasculogenesis in mouse ES cells through the activation of VegfA^34^. These findings validate our experimental design and the robustness of the list of Etv6’s direct target genes.

ETV6 and its Drosophila orthologue, Yan, are well-established transcriptional repressors^14,15,24,41^. Surprisingly, in Xenopus somites, two thirds (303/482) of Etv6 direct target genes were transcriptionally activated. Therefore, during the early stages of specification of the HSC lineage, Etv6 acts both as a transcriptional activator and a repressor. As detailed below, molecular and functional examination of Etv6 target genes revealed that Etv6’s dual function is required for the regulation of vegfa expression.

Members of the ETS family of TFs are defined by a highly conserved ETS domain that recognizes a core sequence 5′-GGA(A/T)-3′ motif within the context of a 9- to 10-bp DNA sequence^42^. As all ETS TFs bind to the same motif, additional mechanisms regulating the selection of specific transcriptional targets within biological contexts are required. It has been suggested that high affinity ETS motifs found in the promoter of housekeeping genes can be bound by any ETS TF whereas lower-affinity ETS binding sites found in tissue-specific promoters and only bound by a subset of ETS TFs are flanked by binding sites of other TFs^43,44^. Cooperative binding with other TFs in sequences with composite binding sites results in a higher affinity and stable binding to DNA, and in synergistic repression or activation of specific target genes^45,46^.

Here, de novo motif analysis of the 498 peaks associated with the promoters of genes normally activated or repressed by Etv6 showed (i) high prevalence of Klf/Sp binding sequences in both sets of targets and (ii) absence of Ets-binding motif enrichment in the peaks associated with genes activated by Etv6. This suggested that members of the Klf/Sp family are required for both activating and repressive transcriptional activities of Etv6 and that Etv6 is recruited to DNA through distinct mechanisms when activating or repressing gene expression.

Strikingly, klf4 was one of the target genes activated by Etv6 in somites. KLF4 is a multifunctional TF which is critical for pluripotency, tissue homeostasis, stemness, maintenance of cancer stem cells and normal hematopoiesis^47^, and has previously been implicated in the expression of VegfA in human and mouse cells both positively and negatively. KLF4 binds the VegfA promoter and represses Vegfa expression during EMT in mouse mammary gland cells^48^, whereas, in human retinal microvascular endothelial cells and HUVECs, it promotes angiogenesis by transcriptionally activating VegfA^27,28^. Here, we show that Klf4 not only activates vegfa expression through binding to conserved Klf/Sp motifs but also recruits Etv6 to the vegfa promoter that, as expected for a gene activated by Etv6, is devoid of Ets-binding sites in the Etv6 peak region. This suggested direct interaction between the two proteins. In support of this, it has been shown, in overexpression experiments with tagged proteins, that the ETS protein ERG co-immunoprecipitates with KLF2^49^. Similarly, in situ proximity ligation assays have demonstrated that KLF4 and the phosphorylated form of the ETS TF, ELK1, physically interact in human coronary artery cells^50^. Moreover, analysis of EHF (an ETS TF structurally related to ETV6) ChIP-seq peaks indicates that a significant number of peaks contained no ETS-binding motifs and many of them harbored Klf4/5 binding motifs only^51^. Interestingly, EHF is involved in the maintenance of corneal transparency through the repression of angiogenic factors such as VegfA^51^. In conclusion, the mechanistic relationship between Etv6 and Klf4 in regulating vegfa expression is compatible with a coherent feed-forward loop, a regulatory circuit where two pathways downstream of a TF control expression of the same gene: Etv6 activates klf4, Klf4 binds and activates vegfa and Etv6 binds vegfa^52,53^ (Figure 6). How Etv6 recruitment to vegfa promoter is functionally integrated to Klf4 activity in the control of vegfa expression remains to be fully explored. Depending on whether both Etv6 and Klf4, or only one of these two regulatory inputs are required for vegfa expression, this circuit is predicted to either delay vegfa expression or control vegfa levels^53^.

**Figure 6.**
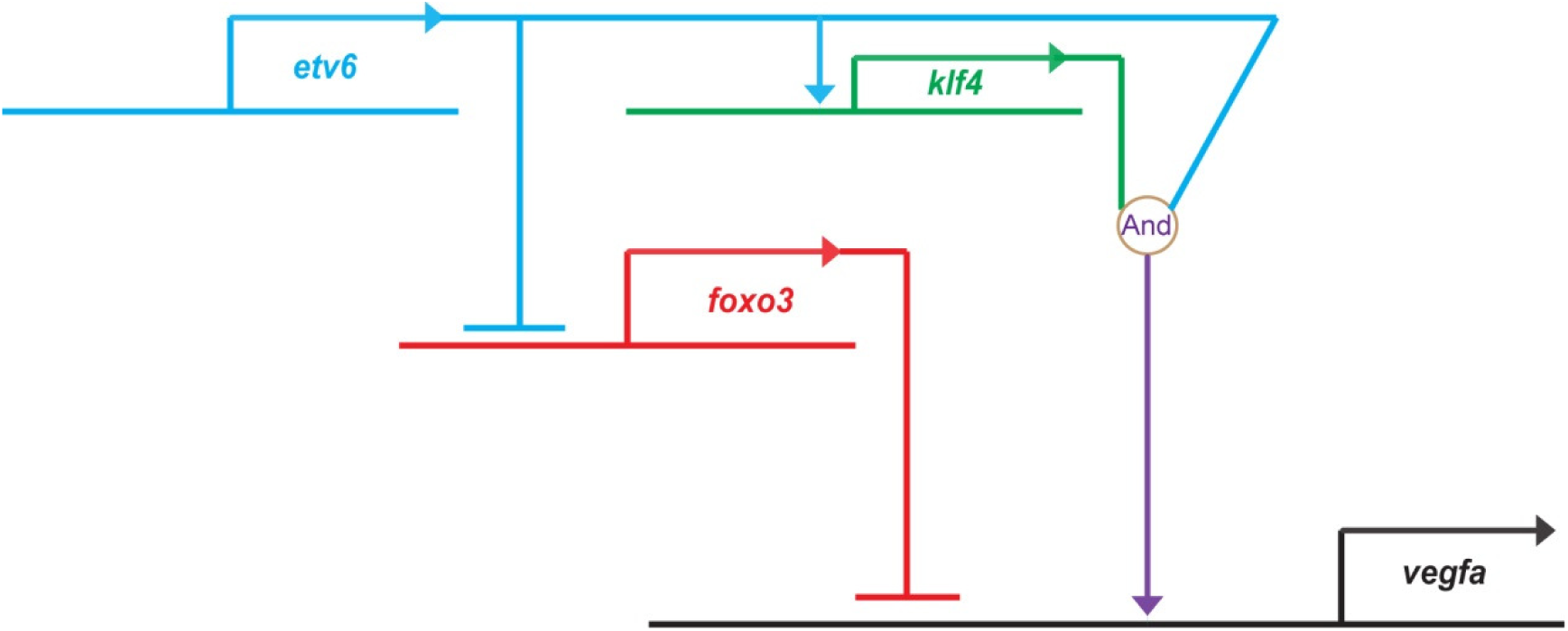
GRN summarizing the mechanisms by which Etv6 regulates the expression of *vegfa* in the somites. Etv6 positively regulates vegfa expression in the somites through multiple mechanisms. It represses the transcription of foxo3, a repressor of vegfa expression. In parallel, it activates the expression of klf4, an activator of vegfa. Finally, Klf4 is required for Etv6 recruitment to the vegfa promoter. In conclusion, a network of positive and negative inputs from Etv6 is required for transcriptional activation of vegfa in the somites.

In addition to activating an activator of vegfa, Etv6 achieves vegfa activation by repressing a transcriptional repressor of vegfa, Foxo3. Consistent with Etv6 direct DNA-binding repressive functions^41^, and in agreement with the distinct features of Etv6-bound activated and repressed promoters, the foxo3 promoter contains an Ets-binding motif. In humans, FOXO3 harbours tumor suppressor activity as it represses processes such as VEGFA-driven tumor angiogenesis^54^. As an example, in breast cancer, FOXO3 activation correlates with VEGFA downregulation^25^. Mechanistically, in human cells, FOXO3 represses VegfA expression by binding to a Forkhead response element in the VegfA promoter^25^. We have confirmed that the Xenopus vegfa promoter contains a conserved Forkhead binding motif (5′-GTAAACA-3′)^39^ and demonstrated that Foxo3 is a repressor of vegfa in the somites as (i) over-expression of this TF results in vegfa repression and (ii) down-regulation of foxo3 in etv6-deficient embryos rescues vegfa expression. We propose that this double negative gate^55^ (Etv6 repressing Foxo3 that represses vegfa), together with the feed-forward loop described above, unlocks the endothelial cell fate in the lateral plate mesoderm through tight control of vegfa expression (Figure 6).

Gene repression by ETV6 and Yan is mediated by the pointed (SAM) and linker domains. These domains can repress transcription independently through different mechanisms. The linker domain represses transcription by complexing with co-repressors which recruit histone deacetylases (HDACs)^14,24,56^. The pointed domain represses transcription by a mechanism that does not involve co-repressors recruiting HDACs^14^. Additionally, the pointed domain has the capacity to form homo- and hetero-typic oligomers and it has been proposed that polymerization of ETV6 could facilitate the spreading of transcriptional repression complexes along long stretches of DNA^16^. However, the formation of ETV6 polymers has not yet been demonstrated in vivo. We found a total of 9,128 Etv6 ChIP-seq peaks in our analysis but only 102 of them were >2 kb in size (Supplementary Table 1) and none were associated with DEGs. Therefore, although long stretches of DNA occupancy are present in the genome, Etv6 polymerization does not mediate gene repression in Xenopus somites.

In conclusion, our investigation of the functions of Foxo3 and Klf4 in vegfa expression in the somites unveils the foundations of a complex Etv6 gene regulatory network (GRN). It indicates that Etv6 has the capacity to regulate vegfa expression through the deployment of pathways involving both positive and negative regulators of gene expression and begins to unveil the complexity of vegfa regulation during hematovascular development. This work also further strengthens the parallel between oncogenic and developmental processes^13^. We propose that this Etv6-vegfa GRN is conserved through vertebrate evolution, from Xenopus to human, and that Etv6 uses, depending on tissue and context, particular components and/or branches of this network to ensure the correct levels of vegfa expression. Further investigation will establish whether Klf4, or additional members of the Klf/Sp family of TFs, are required for the regulation of some of the other Etv6 targets identified in our study and known to regulate Vegfa in the tumorigenic processes. If so, studying these molecular mechanisms could lead to new strategies for the treatment of pathological processes driven by the Etv6-vegfa GRN.

## Methods

### Embryo manipulation

Animal experiments were performed according to UK Home Office regulations under the appropriate project licence and approval of the University of Oxford Animal Welfare and Ethical Review Body. Xenopus laevis embryos were obtained, cultured and injected as previously described^57^. The somites of stage 22 embryos were manually dissected in 0.35x MMR (1xMarc’s Modified Ringer: 100 mM NaCl, 2 mM KCl, 1 mM MgCl_2_, 2 mM CaCl_2_, 5 mM HEPES, pH 7.5. Prepare a 10X stock, and adjust pH to 7.5) using forceps, these explants also contained the hypochord, neural tube, notochord, and some dorsal endoderm. Whole-mount in situ hybridization (WISH) was performed as previously described^57^. Before photography, embryos were cleared in benzylbenzoate:benzyl alcohol (2:1). For probe details see Supplementary Table 8.

Morpholino antisense oligonucleotide (MO), TALEN and mRNA for injection MOs were obtained from GeneTools LLC (Corvallis, OR). Etv6 MO targeting both etv6.L and etv6.S was previously published^12^. A klf4 MO targeting both klf4.L and klf4.S was designed (Figure 4e). As single MO blocking both foxo3.L and foxo3.S could not be designed (Supplementary Figure 3a), two MOs, one targeting foxo3.L (foxo3.L MO) and the other targeting foxo3.S (foxo3.S MO), were co-injected in a 1:1 ratio in order to generate foxo3-deficient embryos (Supplementary Figure 3c). Every MO was titrated in order to determine the optimal concentration for embryo injection. MO sequences are as indicated in Supplementary Table 9.

A two-step Golden Gate assembly method using the Golden Gate TALEN and TAL effector kit 2.0 (Addgene) was used to construct the TALEN plasmids containing the homodimer-type FokI nuclease domain^58^. TALEN sequences were designed using the online design tool, Mojo Hand (http://www.talendesign.org/). TALEN mRNA for injection was generated by linearizing the plasmids with NotI and transcribing with SP6 RNA polymerase using the mMESSAGE mMACHINE Kit (Ambion). TALEN mRNAs were injected at the 1-cell stage of development. To determine the mutagenesis caused by the TALENs, DNA was extracted from the somites of stage 22 TALEN-injected embryos and the genomic DNA fragment containing the TALEN binding sites was amplified and Sanger sequenced (Figure 4f, Supplementary Figure 8).

To make mRNA for injection, the full length coding sequence for X. laevis etv6 and foxo3, including the sequences targeted by the MOs (Supplementary Table 10), were sub-cloned into pBUT3-HA vector (pBUT3-etv6-HA, pBUT3-foxo3-HA). Etv6-HA and foxo3-HA mRNAs were synthesized using mMESSAGE mMACHINE T3 Kit (Ambion).

### Antibodies and Western blot

Polyclonal anti X. laevis Etv6 antibodies were generated in collaboration with NovoPro Bioscience Inc. (Shanghai, China). In brief, the amino acid sequence of X. laevis Etv6 (GeneBank number EU760352) was analyzed using a proprietary algorithm, NovoFocus™ antigen design, to identify epitopes with good hydrophilicity, surface probability and high antigenic index. Based on this information, three epitopes with little or no conservation with other proteins, including ETS TFs, were selected (Supplementary Figure 1a). Short peptides corresponding to these epitopes were synthesized, conjugated to keyhole limpet hemocyanin (KLH) and used in immunizations at a purity ≥90%. Two rabbits were immunized with each peptide and six polyclonal antibodies (Etv6-1a, -1b, -2a, -2b, -3a, -3b) were affinity-purified after 5-6 rounds of immunization. Each antibody had an ELISA titer ≥1:50,000 against the peptide antigen. The specificity of the antibodies was verified by Western Blot analyses.

X. laevis Klf4 protein was detected using an anti-Klf4 antibody (Abcam, ab106629) at a 1:1,000 dilution. HA-tagged protein was detected with anti-HA antibody (Santa Cruz Biotechnology, HA-probe (Y-11)-G) at a 1:500 dilution. Histone H3 protein was detected using anti-Histone H3 (Abcam, ab1791) antibody at a 1:100,000 dilution and β-Actin was detected with anti-β-Actin antibody (Anaspec, AS-55339) at a 1:200 dilution. Clean-blot IP detection kit (Thermo Fisher Scientific, Catalog number: 21232) was used as a secondary antibody in the Western blot for the immunoprecipitation experiments and was used at a 1:200 dilution. Protein extraction and western blot analysis were performed as previously described^59^.

### ChIP-seq

Etv6 ChIP-seq was performed as previously described^60,61^ except that Crosslinker EGS (ethylene glycol bis(succinimidyl succinate)) was added into cell homogenates before fixation and that sonication was performed using Covaris S220. Immunoprecipitated DNA and input DNA were quantified using Qubit Fluorometer and sequencing libraries were constructed with 1ng of DNA using NEBNext Ultra DNA Library Prep Kit (NEB#E7370) for Illumina sequencing. DNA was sequenced (∼ 15-38 million paired reads/library, 2×40 bp read length) using NextSeq 500. Sequencing was performed on three independent biological replicates with corresponding inputs as control.

Raw sequence reads were checked for base quality, trimmed and filtered to exclude adapters using Trimmomatic (Version 0.32)^62^, and then mapped to the X. laevis V9.1^38^ with BWA version 0.7.12. Peaks were analyzed using MACS2 peak calling software^63^ at default thresholds, with the input samples as control for each replicate. Peaks consistent between replicates were identified using DiffBind (Bioconductor package)^64^, those located in TSS regions were identified using Homer software and used to perform TFs binding site analysis with Homer software (findMotifsGenome.pl)^65^. For the TFs binding site prediction in the vegfa and foxo3 promoter region, DNA sequences under the Etv6 peak were analyzed using Jaspar^40^. ChIP-seq data was visualized on the Integrative Genome Viewer (IGV).

### RNA-seq

Total RNA was extracted from somites dissected from 20 WT or Etv6-deficient embryos at stage 22. Triple biological replicates were generated. Indexed libraries were constructed with 1μg of total RNA using the KAPA Stranded RNA-seq Kit with RoboErase (KK8483) and NEBNext Mutiplex Oligos (NEB#E7500S) for Illumina sequencing. DNA was sequenced (∼60-105 million paired reads/library, 2×75 bp read length) using NextSeq 500.

Sequenced reads were checked for base qualities, trimmed and filtered to exclude adapters using Trimmomatic (Version 0.32)^62^ and then mapped to the X. laevis V9.1^38^ using STAR^66^ with default parameters. Aligned read features were counted using Subread tool: featureCounts method (version 1.4.5-p1). Differential gene expression analysis was carried out using EdgR (Bioconductor Package)^67^.

An ANOVA-like test analysis^68^ was performed between WT and MO samples, using the generalized linear model^69^ followed by Estimates “Dispersion”, fitted with “negative binomial model” and estimates “Generalized linear model likelihood ratio”. Genes with a False Discovery Rate (FDR) above 0.05 were filtered out. The functional annotation (gene ontology analysis) of Etv6-regulated transcriptome was performed using DAVID Bioinformatics Resources^70^.

### qPCR

qPCR was performed using Fast SYBR Green Master Mix (Thermo Fisher Scientific, Catalog number:4385612) and StepOne Real-Time PCR system. For RT-qPCR, cDNA was made from 1μg of total RNA and relative expression levels of each gene were calculated and then normalized to odc1 gene. Details of primer sequences for RT-qPCR and ChIP-qPCR are indicated in Supplementary Tables 11 and 12, respectively.

### Statistical analysis

In RT-qPCR and ChIP-qPCR experiments, error bars represent Standard Error of Mean (±SEM). The data shown summarize the results of three biological replicates. Two-tailed Student’s t-test was performed (*p < 0.05; **p < 0.01; ***p < 0.001).

WISH images and numbers shown in Figures are from one experiment and are representative of three biological replicates.

No statistical methods were used to predetermine sample size.

No data or animal have been excluded in this study.

Xenopus laevis embryos were allocated to experimental groups on the basis of different treatments and randomized within the given group. Investigators were not blinded to the group allocation during experiments and outcome assessment.

### Data availability

ChIP-seq and RNA-seq datasets that support the findings of this study have been deposited online in the Gene Expression Omnibus (GEO) under accession number GSE115225. Data can be accessed at https://www.ncbi.nlm.nih.gov/geo/query/acc.cgi?acc=GSE115225 using token mfwtwaeqdjcxbad. Raw pictures for Figure 4e and Supplementary Figures 1b-c, 3b, 8b have been provided in Supplementary Figure 9. All other data that support the findings of this study are available from the corresponding authors upon request.

## Acknowledgement

This work was supported by the UK Medical Research Council (MRC, MC_UU_12009/9) and Biotechnology and Biological Sciences Research Council (BBSRC, BB/M001938/1).

## Author contributions

L.L., R.P., A.C., C.P. designed the experiments. L.L. performed all experiments, prepared the figures and wrote the manuscript. A.C. designed and generated TALEN and designed peptides for ETV6 antibodies production. R.R. performed bioinformatics analyses. R.P., A.C. & C.P. analyzed the data and revised the manuscript. All authors read and approved the final manuscript.

